# Vezatin is required for the retrograde axonal transport of endosomes in *Drosophila* and zebrafish

**DOI:** 10.1101/2020.02.09.940890

**Authors:** Michael A. Spinner, Katherine Pinter, Catherine M. Drerup, Tory G. Herman

**Author notes:** To whom correspondence should be addressed Tory Herman, Ph.D., Institute of Molecular Biology, 1370 Franklin Blvd, University of Oregon, Eugene, OR 97403.

## Abstract

Active transport of organelles within axons is critical for neuronal health. Retrograde axonal transport, in particular, relays neurotrophic signals received by axon terminals to the nucleus and circulates new material among *en passant* synapses. The single retrograde motor, cytoplasmic dynein, is linked to diverse cargos by adaptors that promote dynein motility. Here we identify Vezatin as a new, cargo-specific regulator of retrograde axonal transport. Loss-of-function mutations in the *Drosophila vezatin-like* (*vezl*) gene prevent signaling endosomes containing activated BMP receptors from initiating transport out of motor neuron terminal boutons. *vezl* loss also decreases the transport of endosomes and dense core vesicles (DCVs) within axon shafts. While vertebrate Vezatin (Vezt) has not previously been implicated in axonal transport, we show that *vezt* loss in zebrafish impairs the retrograde movement of late endosomes, causing their accumulation in axon terminals. Our work establishes a new, conserved, cargo-specific role for Vezatin proteins in axonal transport.

## INTRODUCTION

Neurons mediate cognition and behavior by forming networks that can have complex connectivity and span considerable distances. As a consequence, neurons have elongated processes along which passive diffusion is inefficient and materials must instead be actively transported (recently reviewed in Guedes-Dias and Holzbaur, 2019). Within axons, cargoes move along uniformly polarized microtubules by binding to active kinesin or cytoplasmic dynein motors: the former move associated cargo in the anterograde direction, toward microtubule plus ends and the axon terminal, and the latter move cargo in the retrograde direction, back to the soma (Guedes-Dias and Holzbaur, 2019). Bidirectional axonal transport is essential for the function and survival of neurons, and abnormalities in this process are an early hallmark of neurodegenerative diseases such as Huntington’s, Alzheimer’s, and Parkinson’s diseases (Guo et al., 2019).

How axonal transport is regulated remains a central question. While multiple kinesin family members transport the cargos that move toward axon terminals, a single dynein is responsible for the majority of retrograde axonal transport. Dynein’s ability to bind cargo and microtubules is enhanced by its cofactor, dynactin (reviewed in Reck-Peterson et al., 2018). However, a variety of cargo-specific regulators are additionally required (Reck-Peterson et al., 2018). Some link cargoes to dynein-dynactin complexes, some promote the motility of cargo-bound dynein-dynactin, and some do both. While many such regulators have been identified, others undoubtedly remain to be discovered.

The *Drosophila* larval neuromuscular junction (NMJ) is a powerful system in which to identify and analyze genes required for neural development and function, including genes required for axonal transport (Neisch et al., 2016). Each motor neuron forms a stereotyped pattern of branched chains of synaptic boutons on its target muscle and adds new boutons in response to a muscle-secreted Bone Morphogenetic Protein (BMP) signal during larval growth (reviewed in Deshpande and Rodal, 2016). Within larval motor neuron axons, two broad types of cargoes are retrogradely transported: (1) those, such as signaling endosomes containing activated BMP receptors, that originate in the boutons themselves and are transported through the NMJ and axon shaft to the soma (Lloyd et al., 2012; Smith et al., 2012); and (2) those, such as dense core vesicles (DCVs), that originate in the soma, are anterogradely transported to the terminal boutons of each NMJ, but require retrograde transport within the NMJ itself in order to be distributed among the other, *en passant*, boutons (Wong et al., 2012). Both types of cargo accumulate in terminal boutons when dynactin is disrupted by loss of its core subunit p150^Glued^ (Lloyd et al., 2012).

Forward genetic screens in invertebrates continue to uncover new relationships between molecules and phenotypes. We identified a loss-of-function mutation in a previously uncharacterized gene, *CG7705*, that results in enlarged terminal boutons containing accumulated retrograde cargos. CG7705 is most closely related to vertebrate Vezatin (Vezt) proteins. Using reverse genetics and *in vivo* imaging, we show here that both *CG7705*, which we will refer to as *vezatin-like* (*vezl*), and zebrafish *vezt* are required for the retrograde axonal transport of late endosomes. Our data identify a new, conserved role for Vezatin proteins as cargo-specific regulators of endosome transport in axons.

## RESULTS

### Loss of a Vezatin-like (Vezl) protein alters the morphology of terminal synaptic boutons in adult eye and at the larval NMJ

*Drosophila* R7 photoreceptor neurons are a convenient cell type in which to identify essential genes that are required cell-autonomously for neuronal function. Photoreceptors are not required for viability, and the *GMR-FLP*/MARCM technique can be used to generate mosaic animals in which GFP-labeled R7s are homozygous for randomly mutagenized chromosome arms within otherwise heterozygous animals (Lee and Luo, 1999; Lee et al., 2001). In a screen of EMS-induced mutations, we identified a mutation, *721*, that alters R7 axon terminal morphology. Both wild-type and *721* mutant R7 axons terminate in ellipsoid boutons within the M6 layer of the optic lobe (Figure 1A,B). However, *721* mutant R7 terminals also frequently extend short, thin projections that end in distinctive blobs beyond M6 (Figure 1A,B).

**Figure 1.**
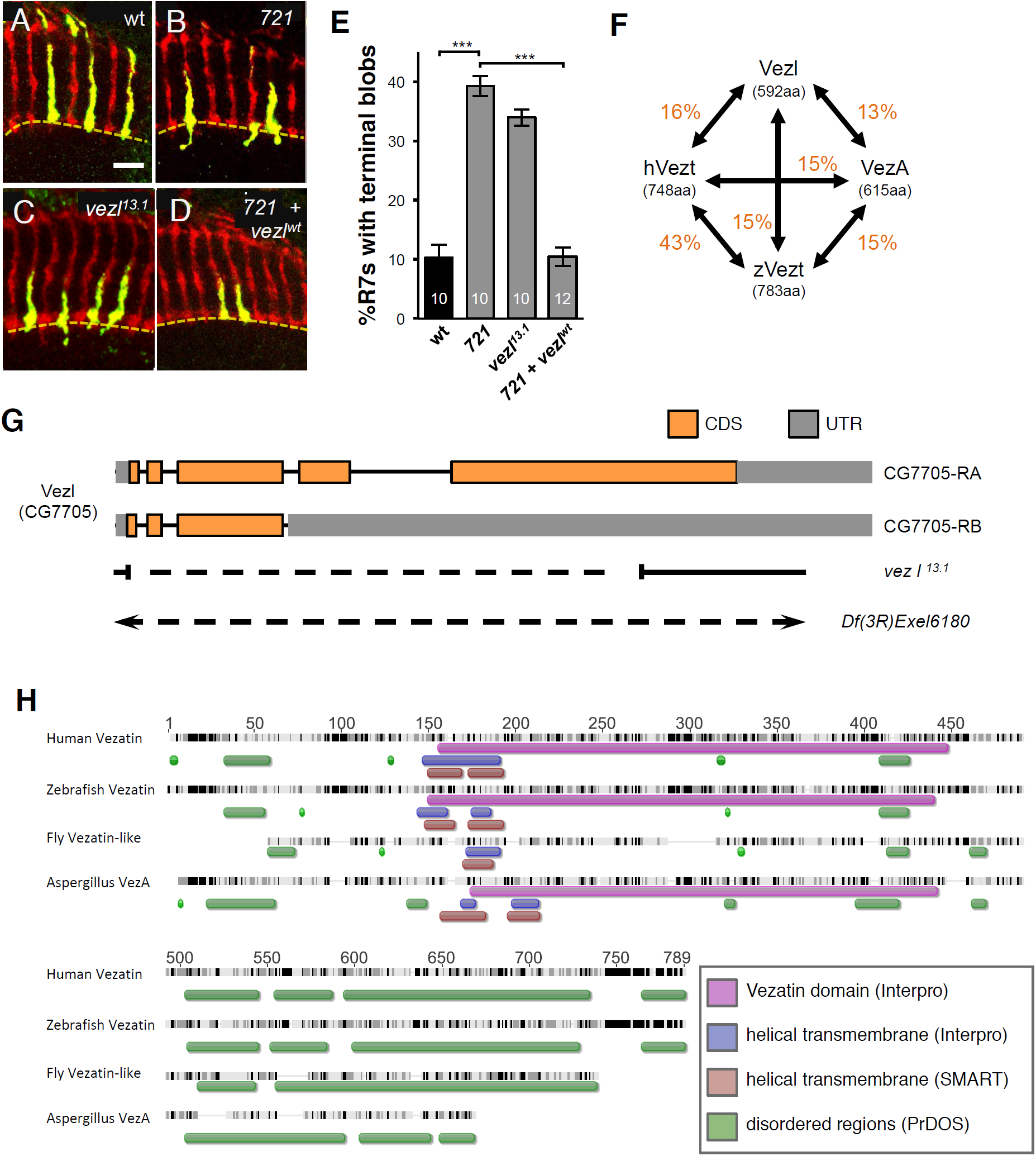
Vezl is required in R7 photoreceptors for normal axon terminal morphology. **A-D**, Adult medullas in which homozygous R7s generated by *GMR-FLP*/MARCM express Syt:GFP (green). All R7 and R8 axons are labeled with anti-Chp (red). The M6 layer of the medulla is indicated by a dashed yellow line. Scale bar is 10 µm. **A**, Wild-type (homozygous *FRT82*) R7 axons terminate in ellipsoid boutons in M6. **B**, The axon terminals of R7s homozygous for the *721* mutation are ellipsoid but frequently have short extensions that end in blobs beyond M6. **C**, R7s homozygous for *13.1*, which deletes most of the *CG7705/vezl* open reading frame, have the same axon terminal defect as *721* homozygous R7s. **D**, Expression of a *vezl* cDNA in *721* homozygous R7s fully rescues their axon terminal defect. **E**, Quantification of extensions in **A-D** genotypes. \*\*\**p*<0.001, based on two-tailed t-tests. Error bars indicate standard error of the mean (SEM). n values (number of animals) are indicated for each bar. As for all bar graphs, the calculated means, SEMs, and p values are available in Supplemental Table 1. **F**, Amino acid sequence comparisons among fly Vezl, human Vezatin (hVez), *A. nidulans* VezA, and zebrafish Vezatin (zVez). Each value in orange indicate the percentage of amino acid identity between a given pair. **G**, Diagram showing the approximate breakpoints of the *vezl*^*13.1*^ deletion relative to the locations of the *vezl* exons found in the two types of *vezl* mRNA identified, “CG7705-RA” and “CG7705-RB”. The first exon is at the left. The large deficiency *Df(3R)Exel6180* completely removes *vezl* together with 23 additional genes. Amino acid coding sequences (CDS) are in orange and untranslated regions (UTR) are in gray. **H**, Sequence alignment of Vezatin family proteins. Black regions indicate conserved sequence and dark grey indicates >80% conserved sequence (Clustal W). A “Vezatin domain” (purple bar) that is defined by the SMART domain predictor is not present in fly Vezl. Helical transmembrane domains predicted by two different algorithms are indicated by blue (Interpro) and red (SMART) bars. Green bars indicate low complexity regions predicted by PrDOS.

Animals homozygous for the *721*-containing chromosome do not survive to adulthood. Using recombination mapping and complementation tests between the *721*-containing chromosome and molecularly-defined deletions, we found that both the lethality and the R7 defect are linked to a previously uncharacterized gene, *CG7705*. The predicted CG7705 protein has no obvious sequence motifs, and simple BLAST searches identified homologs only in other Drosophilid species. However, Position-Specific Iterative (PSI)-BLAST identified a similarity between CG7705 and vertebrate Vezt proteins (Figure 1F,H; Küssel-Andermann et al., 2000). Despite a low percentage of amino acid sequence identity, CG7705 and Vezts share a predicted overall structure, consisting of central alpha-helical transmembrane domains and N- and C-terminal disordered regions. We therefore refer to the *CG7705* gene as “*vezatin-like*” (“*vezl*”). By searching the Vezatin literature, we found a fungal protein, VezA, which was also named for its resemblance to vertebrate Vezt (Figure 1H; Yao et al., 2015), although the percentage of sequence identity in any pairwise comparison among the three proteins is low (Figure 1F).

To verify that *vezl* loss is responsible for the R7 blobs, we used imprecise excision of a previously identified P element insertion within the first exon of *vezl* to generate a deletion allele, *vezl*^*13.1*^, that is missing most of the predicted *vezl* open reading frame (Figure 1G). *vezl*^*13.1*^ homozygotes die either shortly before or after pupariation, and homozygous *vezl*^*13.1*^ mutant R7 axon terminals within heterozygous animals have *721*-like short extensions that end in blobs (Figure 1C,E). Expression of a *UAS-vezl* transgene constructed from the *vezl* cDNA with the largest open reading frame fully rescues this defect (Figure 1D,E). We conclude that *vezl* is required for normal R7 axon terminal morphology.

The observation that *vezl* loss causes lethality led us to wonder whether *vezl* might be generally required for neural development. To test this, we turned to the larval NMJ. We found that *vezl* mutants have significantly fewer boutons per NMJ than wild-type animals, a defect that is fully rescued by expression of a *UAS-vezl* transgene in motor neurons (Figure 2A-C,I). In addition, Vezl loss specifically alters the morphology of NMJ terminal boutons, leaving all other, i.e. *en passant*, boutons apparently unaffected. *vezl* mutant terminal boutons are abnormally large (Figure 2E,F,J) and stain more intensely with the neuronal membrane marker anti-HRP (Figure 2E,F,K). *vezl* mutant terminal boutons are also more frequently surrounded by smaller “satellite” boutons, which, when present, stain intensely with anti-HRP (Figure 2G,L). Each of these mutant defects is fully rescued by expressing Vezl in motor neurons (Figure 2I-L), although expressing Vezl in wild-type motor neurons also has a gain-of-function effect, an increase in small satellite-like boutons that emerge along the whole length of the NMJ (Figure 2H,L). We conclude that Vezl is required in both R7s and motor neurons for normal terminal bouton morphology.

**Figure 2.**
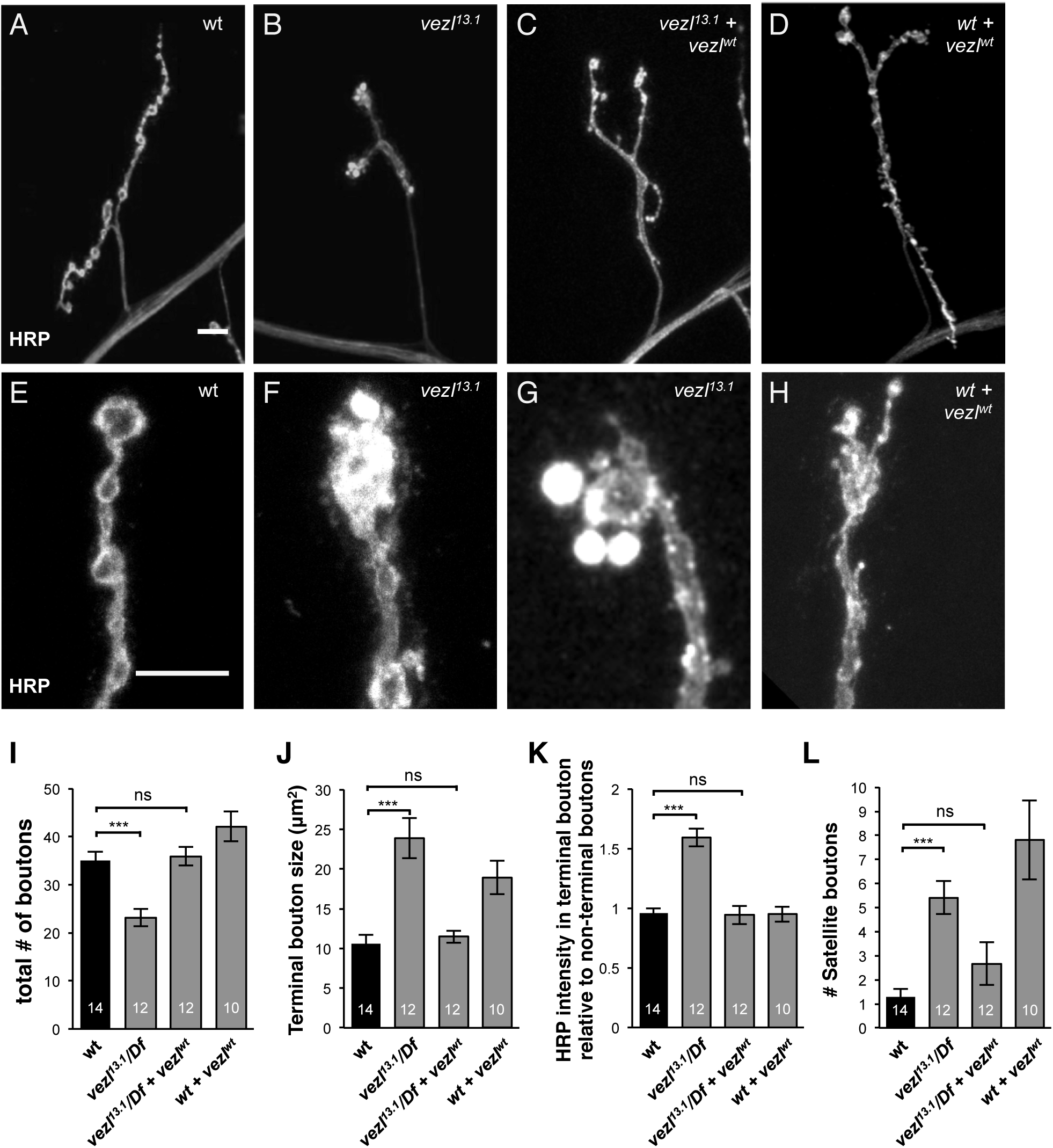
Vezl is required in larval motor neurons for normal NMJ size and terminal bouton morphology. **A-D**, Third-instar larval NMJs at muscle 4 in abdominal segment 3 (A3) stained with anti-HRP (white). Scale bar is 10 µm. **A**, Wild type **B**, *vezl*^*13.1*^*/Df(3R)Exel6180* mutant NMJs have a reduced number of boutons but an increased number of satellite boutons emerging from the terminal bouton **C**, Expression of a *vezl* cDNA in motor neurons of *vezl*^*13.1*^*/Df(3R)Exel6180* mutants restores NMJ size to that of wild type and reduces the number of small, “satellite” boutons emerging from the terminal. **D**, Expression of a *vezl* cDNA in motor neurons of wild-type animals increases the number of satellite boutons throughout the NMJ. **E-H**, Larger magnifications of NMJs stained with anti-HRP (white). Scale bar is 10 µm. **E**, In wild type, the terminal bouton (top) and *en passant* boutons (below) have similar levels of anti-HRP staining. **F**, In *vezl*^*13.1*^*/Df(3R)Exel6180* mutants, the terminal bouton (top left) is often enlarged and contains significantly increased anti-HRP staining. **G**, In *vezl*^*13.1*^*/Df(3R)Exel6180* mutants, the terminal bouton is often surrounded by satellite boutons containing increased anti-HRP staining. **H**, Expression of a *vezl* cDNA in motor neurons of wild-type animals causes an increase in satellite boutons, including those that branch off the terminal bouton (as in this example). **I-L**, Quantifications of the phenotypes mentioned in **A-H**. Expression of a *vezl* cDNA in motor neurons of *vezl*^*13.1*^*/Df(3R)Exel6180* mutants fully rescues these mutant defects. \*\*\**p*<0.001, ns = not significant, based on two-tailed t-tests.

### Vezl loss results in excessive DCVs and endosomes in terminal boutons and disrupts retrograde BMP signaling

Many different manipulations can cause a decrease in bouton number or increase in satellite boutons in fly larval NMJs. However, the only previously reported instance of enlarged terminal boutons with increased anti-HRP staining is that caused by disrupting essential components of the dynein-dynactin complex in motor neurons: knockdown of dynactin or dynein subunits traps dynein-dependent cargoes such as DCVs and endosomes in terminal boutons, causing the latter to expand in size (Lloyd et al., 2012). To test whether Vezl might also be involved in this process, we first examined the localization of retrograde cargoes in fixed samples. We found that Vezl loss, like loss of the essential dynactin subunit p150^Glued^, causes increases in the DCV marker ANF:GFP the early endosome marker YFP:Rab5, and the late endosome marker Rab7:GFP specifically in terminal boutons (Figure 3A-F,I). By contrast, the recycling endosome marker Rab11:GFP, which does not normally undergo retrograde transport, and the synaptic vesicle proteins Synaptotagmin (Syt) and Cysteine String Protein (CSP) do not show such an increase (Figure 3G-I; Supp. Fig. 1A-D). The major active zone (AZ) protein Bruchpilot (Brp) is also not increased in terminal boutons, and Brp puncta remain apposed to postsynaptic glutamate receptors, although their density is decreased (Supp. Fig. 1E-G).

**Figure 3.**
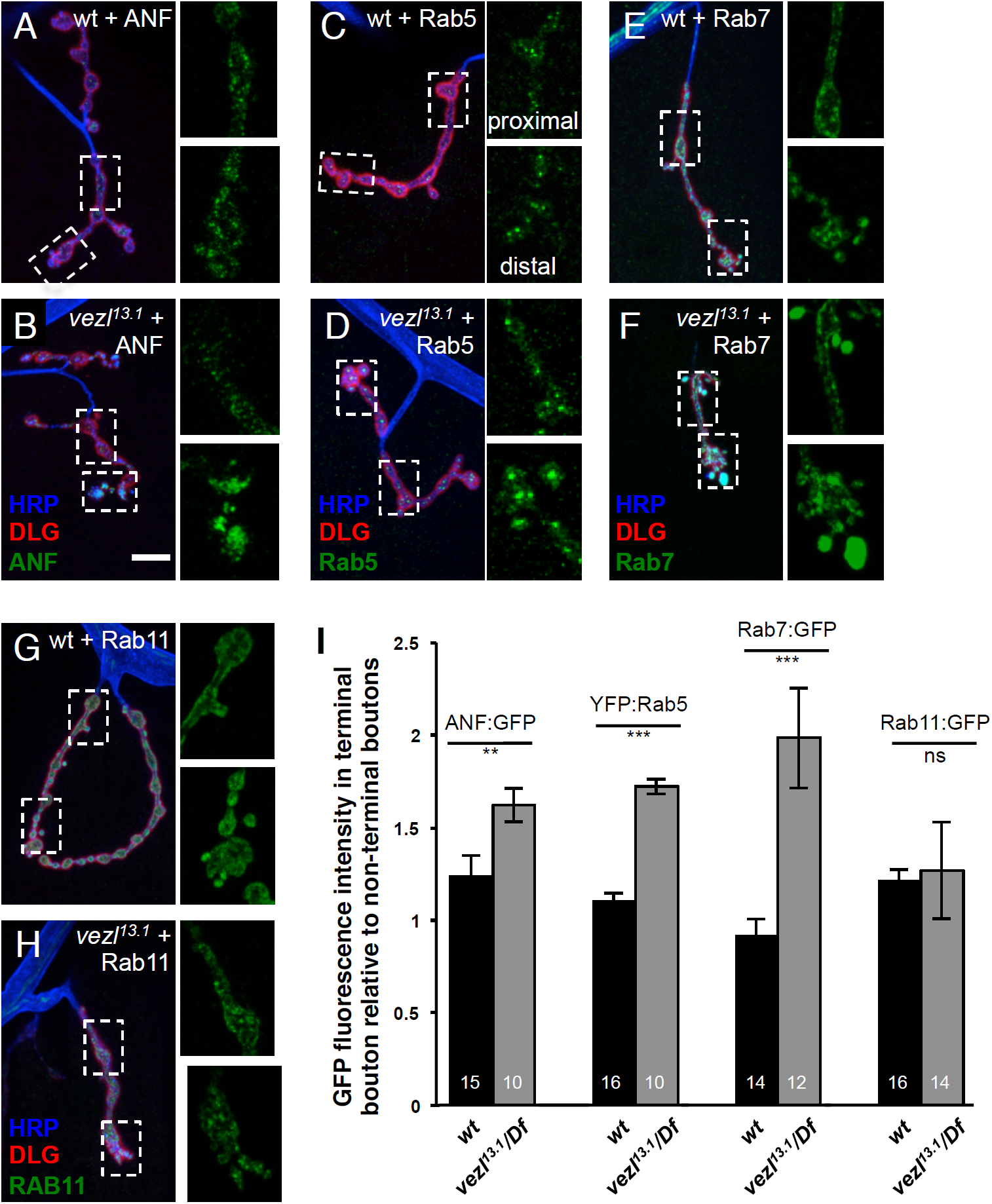
Loss of *vezl* causes an increase in DCVs and endosomes specifically in terminal boutons. **A-H**, Third-instar larval NMJs stained with anti-HRP (blue) and anti-DLG (red). In each panel, dashed boxes indicate the regions containing *en passant* boutons (top) and terminal boutons (bottom) that are displayed with increased magnification in adjacent panels to the right. Scale bar is 10 µm. **A**, In wild-type motor neurons, similar levels of the DCV marker ANF:GFP (green) localize to terminal and *en passant* boutons. **B**, ANF:GFP is increased specifically in terminal boutons in *vezl*^*13.1*^*/Df(3R)Exel6180* mutants. **C**, In wild-type motor neurons, similar levels of the early endosome marker YFP:Rab5 (green) localize to terminal and *en passant* boutons. **D**, YFP:Rab5 is increased specifically in terminal boutons in *vezl*^*13.1*^*/Df(3R)Exel6180* mutants. **E**, In wild-type motor neurons, similar levels of the early-late endosome marker Rab7:GFP (green) localize to terminal and *en passant* boutons. **F**, Rab7:GFP is increased specifically in terminal boutons in *vezl*^*13.1*^*/Df(3R)Exel6180* mutants. **G**, In wild-type motor neurons, similar levels of the recycling endosome marker Rab11:GFP (green) localize to terminal and *en passant* boutons. **H**, The localization of Rab11:GFP in *vezl*^*13.1*^*/Df(3R)Exel6180* mutants resembles that in wild type. **I**, Quantification of GFP intensities in **A-H** genotypes. \*\**p*<0.01, \*\*\**p*<0.001, ns = not significant, based on two-tailed t-tests.

These results implicate Vezl in the initiation of endosomal and DCV transport out of terminal boutons and thereby suggest a potential explanation for the reduced NMJ size in *vezl* mutants. Normal NMJ growth requires motor neurons to transduce a muscle-released BMP signal through the axon to the nucleus (McCabe et al., 2003; reviewed in Deshpande and Rodal, 2016). The BMP signal binds the receptors Wishful thinking (Wit), Saxophone (Sax) and Thick veins (Tkv), which phosphorylate the transcription factor Mothers Against Dpp (MAD). Phosphorylated MAD (pMAD) can then enter the motor neuron nucleus and activate transcription of growth-promoting factors (Marqués et al., 2002; Aberle et a;., 2002; Rawson et al., 2003; McCabe et al., 2003; McCabe et al., 2004). Previous work has shown that pMAD is not retrogradely transported (Smith et al., 2012); instead entry of pMad into the nucleus depends on the retrograde transport of activated BMP receptors that have been trafficked into endosomes (Smith et al., 2012). To determine Vezl’s effect on this pathway, we examined the localization of a tagged form of the BMP receptor Tkv. In fixed samples, Tkv:mCherry normally localizes to puncta distributed throughout the motor neuron axon and boutons (Figure 4A). We found that Vezl loss causes an increase in Tkv:mCherry specifically in terminal boutons (Figure 4D,E), consistent with a defect in its retrograde transport. Previous work has shown that BMP signaling also acts locally, phosphorylating MAD within synaptic boutons by a pathway that does not require retrograde transport (O’Connor-Giles et al., 2008; Higashi-Kovtun et al., 2010; Sulkowski et al., 2016). We found that pMAD levels are significantly increased in *vezl* mutant terminal boutons (Figure 4C-E). Together our results indicate that Vezl loss traps activated Tkv in terminal boutons.

**Figure 4.**
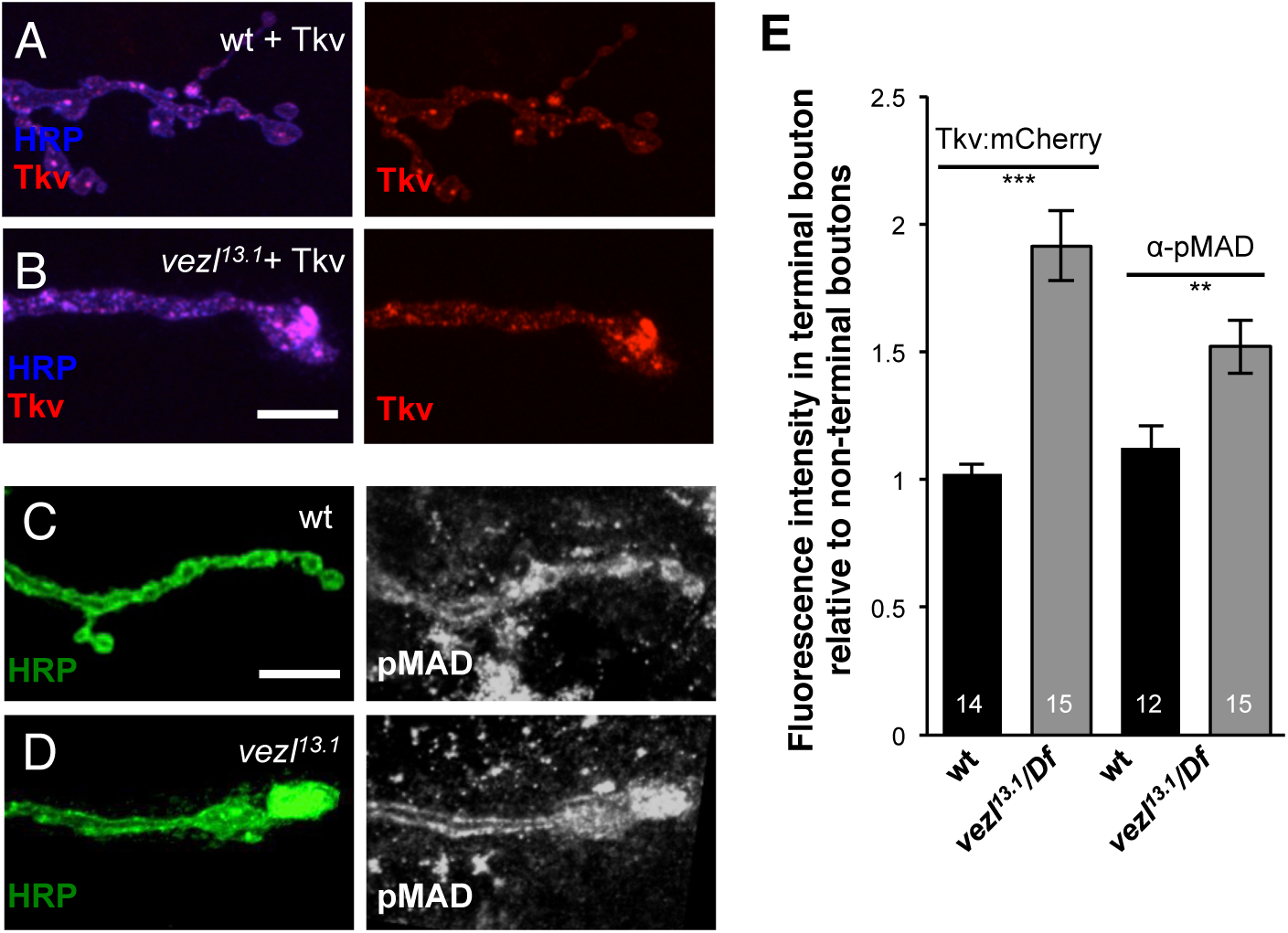
Loss of *vezl* disrupts retrograde BMP signaling. **A,B**, Third-instar larval NMJs stained with anti-HRP (blue) in which motor neurons express Tkv:mCherry (red). Scale bar is 10 µm. **A**, In wild-type motor neurons, similar levels of the BMP type I receptor marker Tkv:mCherry (red) localize to terminal and *en passant* boutons. **B**, Tkv:mCherry is increased specifically in terminal boutons in *vezl*^*13.1*^*/Df(3R)Exel6180* mutants. **C,D**, Third-instar larval NMJs stained with anti-HRP (green) and anti-pMAD (white). **C**, In wild type, similar levels of phosphorylated Mad (pMad), the downstream target of BMP signaling in motor neurons, localize to terminal and *en passant* boutons. **D**, Anti-pMad staining is increased specifically in terminal boutons in *vezl*^*13.1*^*/Df(3R)Exel6180* mutants. **E**, Quantification of staining intensities in **A-D** genotypes. \*\**p*<0.01, \*\*\**p*<0.001, based on two-tailed t-tests.

### Vezl is required for the retrograde movement of Tkv out of terminal boutons

We next directly tested the effect of Vezl loss on the movement of individual vesicles in live animals. We first tracked the movement of Tkv:mCherry. As expected, in wild-type animals significantly more Tkv:mCherry puncta move out of the terminal bouton, i.e. retrogradely, than in the opposite direction (Figure 5A,C). By contrast, in *vezl* mutant animals, almost no Tkv:mCherry puncta move out of terminal boutons (Figure 5B,C); puncta within the terminal boutons move sporadically and non-directionally, reminiscent of Brownian motion (Figure 5B), suggesting that they are not moving along microtubule tracks. We conclude that *vezl* is required for the initiation of Tkv:mCherry retrograde transport out of terminal boutons.

**Figure 5.**
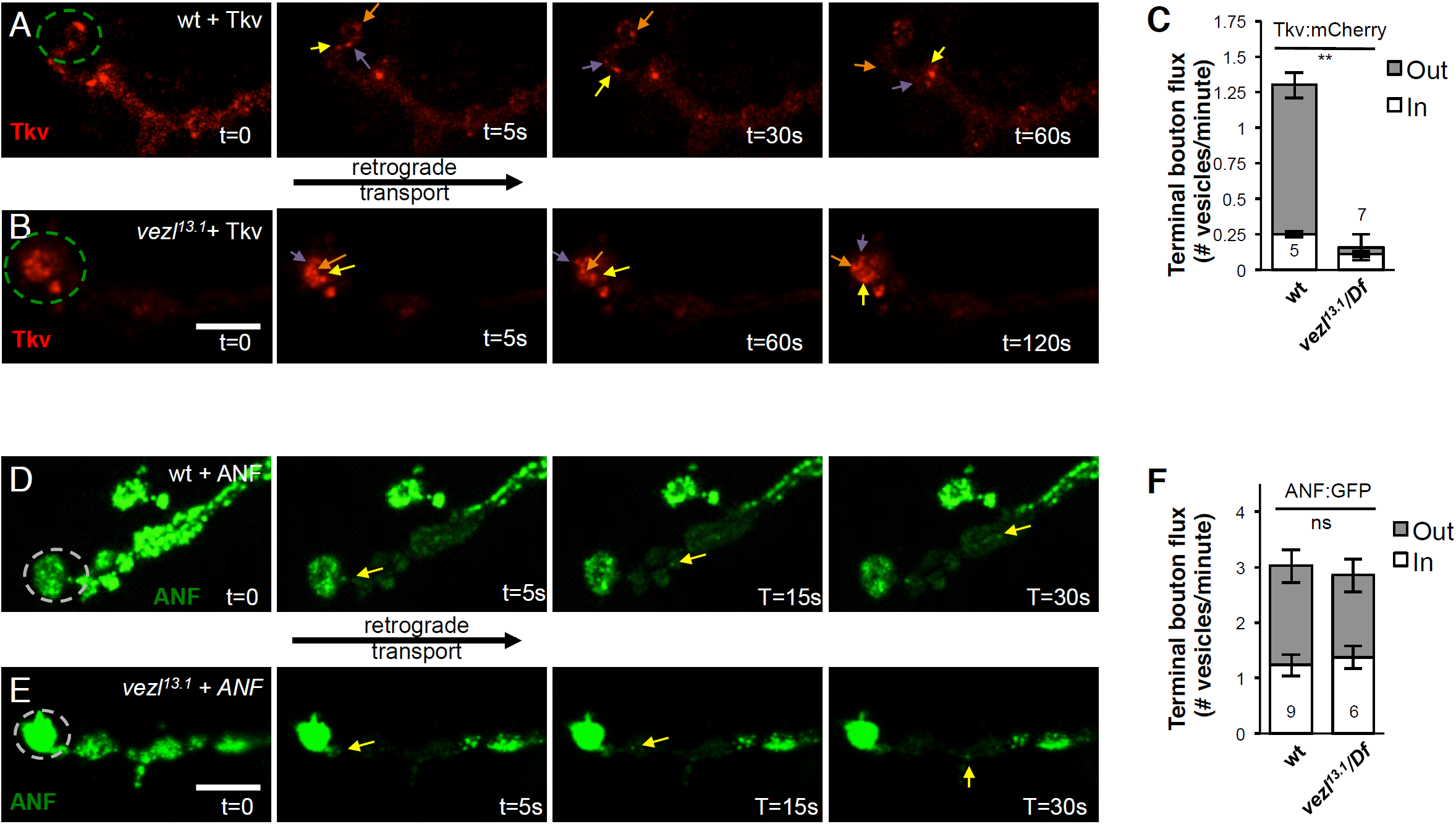
Vezl is required for the retrograde movement of Tkv out of terminal boutons. **A,B,D,E** Timecourses of vesicle movements in live third-instar larvae. Each terminal bouton is on the left (dashed circles) and retrograde transport occurs toward the right. Scale bars are 10 µm. **A**, In wild type, Tkv:mCherry puncta (red; arrows) frequently move retrogradely out of the terminal bouton. **B**, Tkv:mCherry puncta (red; arrows) in *vezl*^*13.1*^*/Df(3R)Exel6180* mutants fail to exit the terminal bouton and instead circulate in an unorganized fashion. Note the extended recording time compared to that in **A**. And note that panel **B** is shown with reduced brightness relative to panel **A** so that individual puncta are more evident. **C**, Quantification of Tkv:mCherry movement. Retrograde movement out of the terminal bouton is significantly decreased in *vezl* mutants. \*\**p*<0.01, based on two-tailed t-tests. **D,E**, Regions adjacent to the terminal bouton were photobleached, allowing the movements of individual ANF:GFP puncta into the bleached region to be monitored. **D**, In wild type, ANF:GFP puncta (green; arrow) frequently move retrogradely out of the terminal bouton. **E**, Movement of ANF:GFP puncta (green; arrow) in *vezl*^*13.1*^*/Df(3R)Exel6180* mutants is similar to that in wild type. **F**, Quantification of ANF:GFP movement. ns = not significant based on two-tailed t-tests.

The possibility remained that *vezl* loss might only affect Tkv:mCherry transport indirectly-perhaps by causing the accumulation of excess HRP-positive membrane. If so, one would expect that most or all retrogradely-transported cargoes would be similarly affected. We therefore next tracked the movement of individual ANF:GFP puncta. Because these puncta are more numerous than Tkv:mCherry and therefore difficult to follow individually, we used fluorescence recovery after photobleaching (FRAP). We photobleached the boutons adjacent to the terminal bouton and measured the number of vesicles that entered the bleached area either by retrograde transport from the terminal bouton or by anterograde transport from more proximal boutons. As expected of cargoes that use retrograde transport to circulate among *en passant* boutons, we found that ANF:GFP puncta do not display the bias toward retrograde movement that we observed for Tkv:mCherry puncta: in wild-type animals, roughly equal numbers of ANF:GFP puncta move out of and toward the terminal bouton (Figure 5D,F). Despite the mild accumulation of ANF:GFP in terminal boutons observed in our fixed images of *vezl* mutants, we found that *vezl* loss does not significantly alter anterograde or retrograde flux at terminal boutons (Figure 5E,F). We conclude that Vezl loss does not prevent all cargoes from exiting the terminal boutons but specifically affects Tkv:mCherry.

### Vezl is enriched in terminal boutons but is also required for normal transport of DCVs and Tkv along axons

Retrograde transport out of terminal boutons is initiated when dynein-cargo complexes are loaded onto microtubules by the dynactin subunit p150^Glued^ (Lloyd et al., 2012; Moughamian and Holzbaur, 2012; Moughamian et al., 2013). Presumably to facilitate this, p150^Glued^ is highly enriched in terminal boutons, unlike dynein itself (Lloyd et al., 2012; Moughamian and Holzbaur, 2012). To test whether Vezl might specifically regulate transport out of terminal boutons, we examined Vezl localization by driving expression of Venus-tagged full-length Vezl in motor neurons of wild-type animals. We found that Vezl:Venus is highly enriched in terminal boutons, consistent with its role in transport initiation (Figure 6A). However, Vezl:Venus is also present at low levels in non-terminal boutons and along axon shafts. We therefore tracked the movement of ANF:GFP and Tkv:mCherry puncta within nerve bundles. In wild type, most such puncta are actively moving (Figure 6B,C,G). We found that *vezl* mutant nerves contain fewer ANF:GFP and Tkv:mCherry puncta than are present in wild type, and a greater proportion of these puncta are stationary (Figure 6D-G). While both anterograde and retrograde flux of each marker is reduced (Figure 6H), retrograde flux is significantly more affected, resulting in puncta that move primarily in the anterograde direction (Figure 6D,E,H). Vezl loss does not alter the velocity of ANF:GFP movement, indicating that motor protein kinetics are unaffected, although we did observe a mild decrease in the anterograde velocity of Tkv:mCherry puncta (Figure 6J). To test the specificity of these effects we also tracked the movement of mitochondria within nerve bundles using Mito:GFP. We found that the proportion of motile Mito:GFP puncta and the direction of their movement are unaffected by Vezl loss (Figure 6F-J). We conclude that Vezl has a cargo-specific role in promoting retrograde transport within axon shafts.

**Figure 6.**
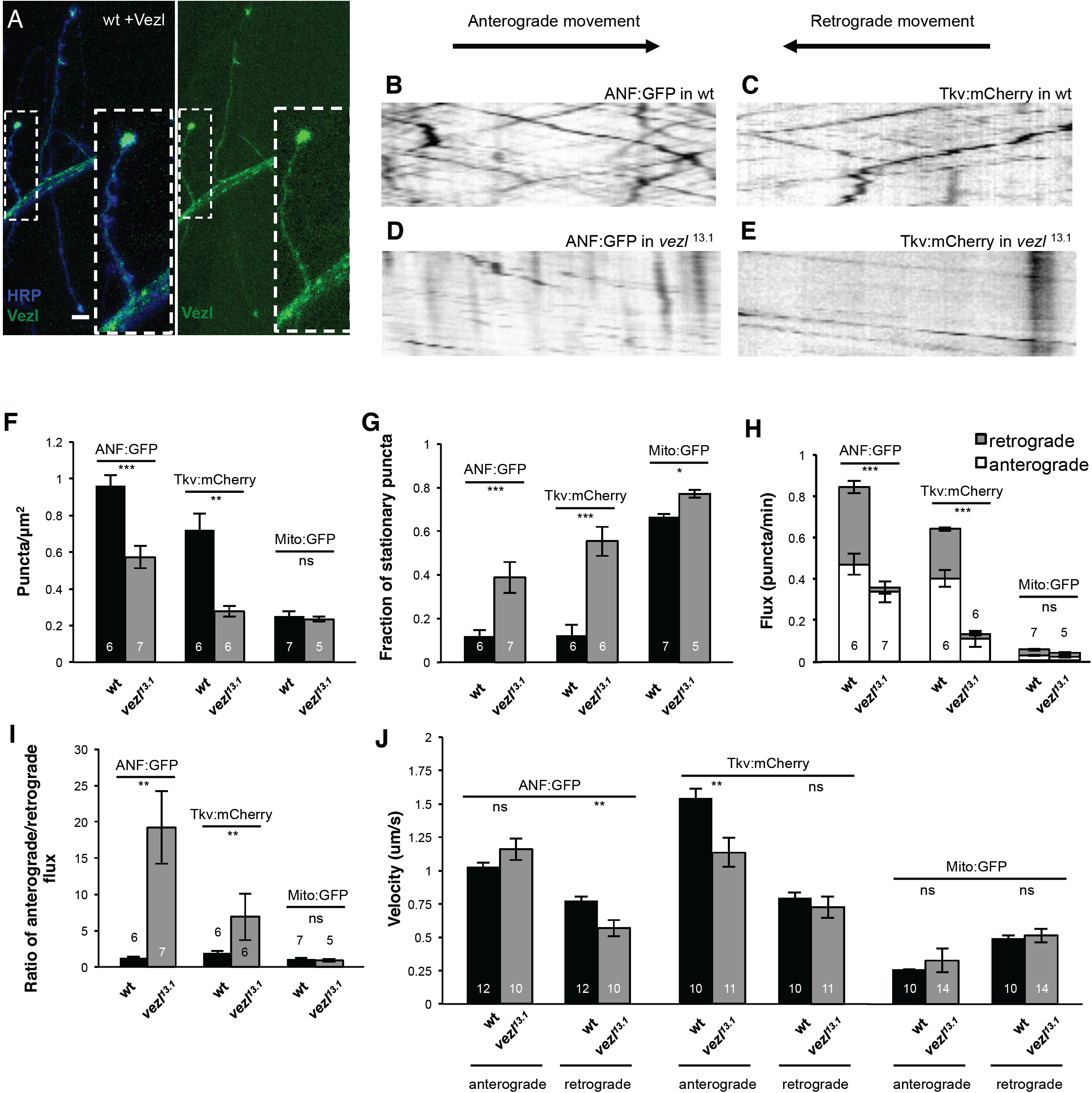
Vezl is enriched in terminal boutons but is also required for normal transport of DCVs and Tkv along axons. **A**, A wild-type third-instar larval NMJ at which Vezl:Venus (green) is expressed in the motor neuron. Anti-HRP is in blue. Vezl:Venus is highest in terminal boutons but also present in en passant boutons and the axon shaft. Scale bar is 10 µm. **B,C,D,E**, Representative kymographs of ANF:GFP (B,D) or Tkv:mCherry (C,E) movement in third instar larval axons. The x-axes represent distance and y-axes represent time. Lines that slope downward from left to right are puncta moving in the anterograde direction; lines that slope downward from right to left are puncta moving in the retrograde direction. Stationary puncta result in vertical lines. In wild type, ANF:GFP (**B**) and Tkv:mCherry (**C**) puncta move in both directions. In *vezl*^*13.1*^*/Df(3R)Exel6180* mutants, retrograde movement of ANF:GFP (**D**) and Tkv:mCherry (**E**) puncta is eliminated. **F-J**, Quantification of ANF:GFP and Tkv:mCherry movement in **B-E** genotypes. Retrograde flux of both ANF:GFP and Tkv:mCherry is signficantly decreased; anterograde flux of Tkv:mCherry is also significantly decreased. \**p*<0.05, \*\**p*<0.01, \*\*\**p*<0.001, ns = not significant, based on two-tailed t-tests.

### Vertebrate Vezatin is required for normal retrograde transport of Rab7-positive endosomes

Vertebrate Vezt has not previously been implicated in axonal transport. To test its potential role in this process, we first examined whether expression of tagged human Vezt (hVez) might rescue fly *vezl* mutant NMJ defects, despite the low amino acid sequence identity between the fly and human proteins. We found that, unlike fly Vezl, hVezt expressed in fly motor neurons is not enriched at terminal boutons and does not restore either NMJ size or terminal bouton morphology in *vezl* mutants (Supplemental Figure 2). We therefore turned to a vertebrate model system, zebrafish, to directly test whether vertebrate Vezt plays a role in retrograde axonal transport. The posterior lateral line (pLL) sensory axons of the larval zebrafish are ideal for analyses of cargo localization and transport in axons, in part because of their considerable length (Drerup and Nechiporuk, 2016). We used CRISPR technology to create a zebrafish line in which the single *vezt* ortholog within the zebrafish genome (Figure 1F,H) is disrupted. In this line, a 5 base pair insertion in exon 3 disrupts the open reading frame, leading to a 40 amino acid insertion and premature stop site.

As in fly, loss of dynein or dynactin in zebrafish causes retrogradely-transported cargoes to accumulate in axon terminals, resulting in characteristic axon terminal swellings (Drerup and Nechiporuk, 2013; Drerup et al., 2017). We found that pLL axon terminals in *vezt* mutant fish have similar large swellings at 4 days post-fertilization (dpf), whereas their wild-type siblings do not (Supplemental Figure 3A-F). To determine whether loss of retrograde cargo movement was responsible for this defect, we first analyzed the localization of dynein and dynactin components in pLL axons by immunofluorescence. We found that excess p150 accumulated in *vezt* mutant terminals as compared to wild-type siblings at 4 dpf (Supplemental Figure 3A,B,G), suggestive of a defect in retrograde transport. Dynein heavy chain (DHC) and mitochondria were normally localized (Supplemental Figure 3C-G).

To directly assess the localization and transport of endosome subtypes, we mosaically expressed mRFP-tagged Rab5, Rab7, and Rab11 in live animals. We found that Rab7:mRFP-positive late endosomes significantly accumulate in pLL axon terminals in *vezt* mutants at 4 dpf, while Rab5- and Rab11-labeled endosomes localize normally (Figure 7A-C,E-G,I-K). We then analyzed the movement of these markers in axons and found that retrograde transport of Rab7:mRFP-positive endosomes is specifically impaired in *vezt* mutants, as assessed by distance traveled (Figure 7H); Rab5-positive early endosome anterograde transport showed a modest decrease that failed to reach significance (Figure 7D), and recycling endosomes moved normally (Figure 7L). We conclude that Vezt is required for the retrograde transport of Rab7-labeled endosomes in vertebrate axons.

**Figure 7.**
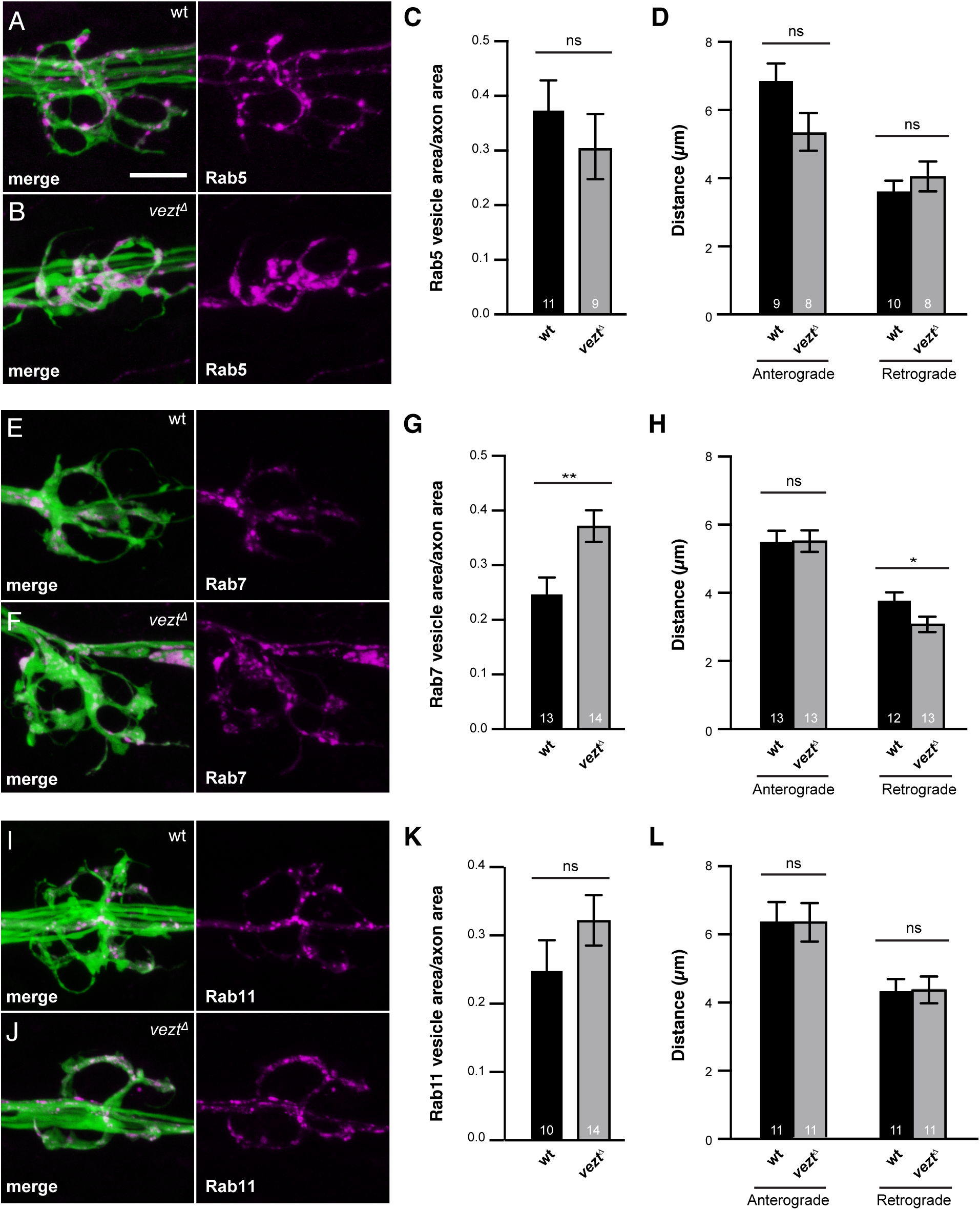
Zebrafish *vezt* mutants have decreased retrograde transport of late endosomes, which accumulate in axon terminals. **A,B,E,F,I,J**, pLL axon terminals marked by cytoplasmic GFP (green) in the *TgBAC(neurod:egfp)*^*nl1*^ transgenic line. Arrowheads point to axon terminal swellings in *vezt* mutants. Scale bar is 10 µm. **A,B**, The localization of Rab5:mRFP (magenta) is similar in wild-type (**A**) and *vezt* mutant animals (**B**). Quantified in **C,D**, which also shows largely normal movement of Rab5:mRFP. **E,F**, By contrast, Rab7:mRFP (magenta) accumulates significantly more in *vezt* mutant axon terminals (**F**) than in wild-type terminals (**E**). Quantified in **G,H**, which also shows a decreased retrograde transport distance in *vezt* mutants. **I-L** Rab11:mRFP localization and transport is unaffected in *vezt* mutants.

## DISCUSSION

Here, we describe the identification of a Vezt-like fly protein, Vezl, that acts as a cargo-specific regulator of retrograde transport. In the absence of Vezl, signaling endosomes accumulate within terminal synaptic boutons and fail to move retrogradely within axon shafts of motor neurons. We find that loss of Vezt from zebrafish similarly impairs the retrograde movement of Rab7-labeled endosomes in sensory axons. This work identifies a new regulator of dynein-dependent endosomal transport in neurons.

While neither fly Vezl nor vertebrate Vezt have previously been implicated in axonal transport, a distantly related *Aspergillus* protein, VezA, is required for the dynein-dependent retrograde transport of endosomes in hyphae (Yao et al., 2015). Tagged VezA is enriched at microtubule plus ends within hyphal tips as Vezl is at terminal boutons, and loss of VezA, Vezl, or Vezt decreases the frequency of retrograde endosome movement, causing endosomes to accumulate at microtubule plus ends (Yao et al., 2015). VezA is thought to act by promoting binding between the endosome-specific cargo adaptor HookA and the dynein-dynactin complex (Yao et al., 2015). Might Vezl and Vezt act similarly?

While the Hook family of cargo adaptors is conserved across species (Xiang et al., 2015; Reck-Peterson et al., 2018), the single Hook protein in fly does not seem to be required for retrograde axonal transport of endosomes. Fly Hook can associate with Rab5- and Rab7-positive endosomes and has some effects on endosome trafficking (Krämer and Phistry, 1999), but *hook* null mutants are viable (Krämer and Phistry, 1999; Sunio et al., 1999; Szatmári et al., 2014), their NMJs do not have the terminal bouton swelling that is characteristic of *vezl* mutants (Narayanan et al., 2000), and their NMJs have more boutons rather than fewer (Narayanan et al., 2000). Nevertheless, the cargo specificity of Vezl - toward endosomes and, to a lesser extent, DCVs but not mitochondria - indicates that, like VezA, Vezl does not alter the general properties of the dynein-dynactin motor or its microtubule substrate. Instead, Vezl likely promotes interactions between cargoes and dynein-dynactin that are required for the initiation of retrograde transport both out of terminal boutons and within axon shafts. Such interactions could be direct or could instead involve another cargo adaptor such as Bicaudal, which shares a similar structure and interaction space on the dynein motor with Hook proteins (McKenney et al., 2014; Schlager et al., 2014; Urnavicius et al., 2018).

Vertebrate Vezt was originally identified based on its association with cell-cell junction proteins, including the cadherin-catenin complex within epithelial cells (Küssel-Andermann et al., 2000; Hyenne et al., 2005), PSD95 within hippocampal dendrites (Danglot et al., 2012), and, more recently, muscle acetylcholine receptor at NMJs (Koppel et al., 2019). Consistent with this localization pattern, Vezt is required for the maturation and maintenance of adherens junctions (Hyenne et al., 2005; Hyenne et al., 2007), dendrites (Sanda et al., 2010; Danglot et al., 2012), and NMJ synapses (Koppel et al., 2019). The precise mechanism by which Vezt has these effects is unclear, but its binding partners include the small GTPase Arf6 (Sanda et al., 2010), which has multiple effects, including on dynein-dependent endosome movements during cytokinesis, exocytosis, and macropinocytosis (Montagnac et al., 2009; Marchesin et al., 2015; Williamson and Donaldson, 2019). It is therefore possible that Vezt’s ability to interact with Arf6 contributes to its regulation of endosome movement in axons. Unlike fly Hook, vertebrate Hooks are well-established as activating adaptors for cargo-bound dynein-dynactin (Reck-Peterson et al., 2018). Zebrafish Vezt may therefore act analogously to VezA and promote binding between the Rab7-specific Hook protein Hook3 and Rab7-endosome-bound dynein-dynactin.

A significant consequence of Vezl loss at fly NMJ is its disruption of neurotrophic BMP signaling. The failure of Tkv to undergo retrograde transport out of terminal boutons provides a simple explanation for the reduced number of boutons at *vezl* mutant NMJs. The observation that pMAD is increased in *vezl* mutant terminal boutons suggests that Vezl loss does not prevent the trafficking of activated receptors into endosomes but instead specifically prevents their transport. We note that overexpressing Vezl in wild-type motor neurons may cause an increase in bouton number (Figure 2I; p=0.0656, two-tailed t test), suggesting that Vezl may be rate-limiting for retrograde BMP signaling.

As we were completing this work, a paper reported the identification of *vezl* mutants in a screen for larvae with defects in cell division: *vezl* loss was found to disrupt the behavior of mitotic and meiotic spindles and their attachment to chromosomes (Graziadio et al., 2018). Despite this defect, *vezl* mutants do not die until they are late larvae or early pupae, presumably because wild-type *vezl* mRNA is supplied by the maternal germline (Fisher et al., 2012). Graziado et al. (2018) did not recognize the similarity of Vezl to vertebrate Vezt or identify the mechanism by which Vezl affects spindle behavior. One possibility is that Vezl regulates dynein-dynactin-dependent movements during cell division as well as in axons. In support of this, Graziadio et al. (2018) observed that CG7705/Vezl loss causes the failure of a dynein-dependent event during mitosis: in *CG7705/vezl* mutants, the dynein-associated protein Zw10 fails to spread along spindle microtubules and instead accumulates at kinetochores. Alternatively, like vertebrate Vezt, Vezl may have functions that are independent of its effects on dynein-dependent transport.

## ACKNOWLEDGMENTS

This work was supported by National Institutes of Health (NIH) grants R03NS100027 (to T.G.H.), T32GM007759 (M.A.S.), and 1ZIAHD008964-02 (to C.M.D). Stocks obtained from the Bloomington *Drosophila* Stock Center (supported by NIH grant P40OD018537) were used in this study, as were DNA constructs obtained from the *Drosophila* Genomics Resource Center (supported by NIH grant 2P40OD010949). The 24B10 and 6D6 antibodies developed by S. Benzer, the 4F3 antibody developed by C. Goodman, the 3H2 antibody developed by K. Zinn, and the nc82 antibody developed by E. Buchner were obtained from the Developmental Studies Hybridoma Bank, which was created by the NICHD of the NIH and is maintained at The University of Iowa.

## MATERIALS AND METHODS

### *Drosophila* genetics

Flies were grown on standard food at 25°C. The *721* mutation was the result of exposing wild-type *FRT82*-containing flies (Bloomington Drosophila Stock Center (BDSC) stock #1359) to ethylmethane sulfonate (EMS) using standard methods (Ashburner, 1989). The *vezl*^*13.1*^ deletion was generated by imprecise excision of a *white+* (*w+*) P element within exon 1 (BDSC#31820). Briefly, *w-*lines generated by mobilizing the P element were tested for their ability to complement *721* for lethality. DNA from those that failed to complement *721* was sequenced with primers flanking the *CG7705* gene region. The *13.1* deletion removed most of *CG7705* without extending beyond it. Homozygous wild-type (*FRT82*), *FRT82 vezl*^*721*^, or *FRT82 vezl*^*13.1*^ R7s were created by *GMR-FLP*-induced mitotic recombination (Lee et al., 2001) and labeled by the MARCM technique (Lee and Luo, 1999) using *actin-Gal4* to drive *UAS-syt-GFP. vezl*^*13.1*^*/Df(3R)Exel6180* homozygous larvae are viable as late third instar larvae but die either shortly before or after pupariation. All NMJ analyses were performed using motile third instar larvae responsive to touch.

The *UAS-Vezl*^*wt*^:*Venus* and *UAS-hVez:Venus* transgenes were generated by cloning a full-length *vezl* cDNA (LP22035; obtained from the Berkeley Drosophila Genome Project) and a full-length hVez cDNA (BC064939, obtained from Dharmacon) into pUAST vectors containing a C-terminal Venus open reading frame (Wang et. al, 2012, Addgene plasmid 35204). Transgenes were sequenced and then injected into embryos containing an attP8 site on the X chromosome (BDSC stock #32233, BestGene).

Flies containing *UAS-Tkv:mCherry* were a generous gift from Thomas Kornberg (Roy et al. 2014). Stocks containing all other transgenes used were obtained from BDSC: *OK371-Gal4* (#26160); *Df(3R)Exel6180* (#7659); *UAS-ANF:GFP* (#7001); *UAS-YFP:Rab5* (#50788); *UAS-Rab7:GFP* (#42705); *UAS-Rab11:GFP* (#8506); and *UAS-mito:GFP* (#8442). Because many transgenes used were located on the X chromosome, we analyzed females only for consistency

### Sequence alignment and domain prediction

The third iteration of PSI-BLAST (https://blast.ncbi.nlm.nih.gov/) Identified vertebrate Vezatins as having sequence similarity to Vezl. Sequence alignments (Figure 1H) were performed with ClustalW2 (Geneious). Domains indicated in Figure 1H were predicted using Interpro (https://www.ebi.ac.uk/interpro/), SMART (http://smart.embl-heidelberg.de/), or PrDOS (http://prdos.hgc.jp/cgi-bin/top.cgi) for disordered domain prediction.

### *Drosophila* dissections and immunostaining

Adult brains were dissected and fixed in 4% paraformaldehyde at room temperature for 20 minutes. Late third-instar larval NMJs were dissected in Schneider’s insect medium (Sigma-Aldrich) and fixed with Bouin’s solution (Sigma-Aldrich) for 5 min. All samples were then stained by standard methods: washed 3× in PBS containing 0.3% Triton-X 100 (PBT), blocked for 30 min in 5% normal goat serum, incubated with primary antibodies at 4°C overnight, washed 3× with PBT, incubated with secondary antibodies at 4°C overnight, washed 3× with PBT, and mounted in Vectashield (Vector Laboratories). We used the following primary antibodies from the Developmental Studies Hybridoma Bank (DSHB): anti-Chp (24B10, at dilution 1:200); anti-Dlg (4F3, 1:250); anti-CSP (6D6, 1:250); anti-Syt (3H2 2D7, 1:250); and anti-Brp (nc82, 1:100). Other primary antibodies used were: rabbit anti-GFP (1:1000; Abcam); fluorescence-conjugated anti-HRP (1:250; Jackson Immuno Labs); rabbit anti-GFP (1:1000; Abcam); rabbit anti-pMAD (1:500, a gift from Carl Heldin; Persson et al. 1998); and rabbit anti-GluRIIC (1:2500, a gift from Aaron DiAntonio; Marrus et al. 2004). All secondary antibodies were goat IgG coupled to Alexa Fluor 488, Alexa Fluor 555, or AlexFluor 633 (1:500; Thermo Fisher).

### Quantification of *Drosophila* R7 and NMJ morphology

For all comparative experiments, animals from each genotype were grown, dissected, and stained in parallel and imaged on the same day under identical microscope settings. Confocal images were collected on a Leica SP2 microscope (with a 63× 1.4 NA oil immersion objective). Maximum-intensity Z projections of confocal stacks were generated and analyzed with ImageJ software (Schindelin et al. 2012; http://fiji.sc/). Within each experiment, the images were randomized, and quantifications were performed blind to genotype. As is standard, we counted boutons based on shape, as visualized by anti-HRP and anti-Dlg staining on muscle 4 on both sides of segment A3 (e.g. Ball et al. 2010). Satellite boutons were identified as smaller, non-chain-forming, individual boutons that emerged from larger boutons on the main NMJ axis (e.g. Marie et al., 2004); they were counted separately and not included in total NMJ bouton counts. The relative sizes and staining intensities of terminal boutons were measured in ImageJ by manually tracing each terminal bouton and its immediate neighbors.

### *Drosophila* live imaging and analysis

Wandering late third-instar larvae were dissected to expose the area of interest in Ca^2+^-free HL3 media on a custom made Sylgard mount, secured with a coverslip, and immediately imaged in a temperature-controlled room during similar times of day. Live imaging was conducted with an Elyra SP.2 camera on a Zeiss LSM 680 using Plan-Apochromat 63x/1.4 oil immersion lens. Videos were recorded with a minimum of two frames per second over the course of 6-10min timespans to ensure the larvae did not die during imaging. We used FRAP to track individual ANF:GFP puncta movement within the NMJ by bleaching a region of interest adjacent to the terminal bouton with a 488nm laser for 60s at high power followed by regular image acquisition. In all cases, the number of puncta that entered or left the terminal bouton over the movie’s timespan was counted manually. Velocities were calculated based on the slopes of kymographs generated by the Multikymograph tool in ImageJ. In axon, the number of puncta were counted and then watched for directional movement over a fixed timeframe of 3 to 5 minutes in order to categorize them as stationary, anterograde-moving, and retrograde-moving. These numbers were normalized to area of the region being imaged.

### Zebrafish husbandry and generation of the *vezt* mutant line

Adult zebrafish were housed at 28°C and bred using established protocols (Westerfield, 1993) in accordance with NICHD/IACUC protocol Drerup 18.008. Embryos and larvae were housed at 28°C in embryo media and staged by established methods (Kimmel, Ballard, Kimmel, Ullmann & Schilling, 1995). The *TgBAC(neurod:egfp)*^*nl1*^ transgenic line was used to label neurons with cytoplasmic GFP (Obholzer et al 2008).

The *vezt* mutant line was generated using CRISPR/Cas9 mutagenesis (Burger et al. 2016). Briefly, 500ng Cas9 protein (Integrated DNA Technologies) was coinjected with ∼200pg gRNA targeted against exon 3 of the zebrafish *vzt* ortholog (GTGGCGCTGTCTCGACGCAG). Resulting F0 animals were raised and crossed to wild-type animals to generate F1s.

Sequencing of the F1 larvae revealed an F0 carrier of a 5 base pair insertion in exon 3 of *vezt*, which leads to insertion of 40 amino acids followed by a premature stop site. Fragment length analysis of PCR products flanking this region were used to genotype (Carrington et al. 2015). The *vezt* mutation is homozygous lethal, and the axon terminal swelling phenotype it causes is recessive.

### Immunolabeling and fluorescence intensity quantification in zebrafish

Homozygous embryos were generated by *vezt* heterozygous crosses. Larvae were raised to 4 dpf, fixed in 4% paraformaldehyde, and stained as in Drerup and Nechiporuk (2013). Antibodies used were: Anti-Dync1h1 (Proteintech #12345), Anti-p150 (BD transduction laboratories #610473), Anti-Cytochrome C (BD Pharma #556432), Anti-GFP (Aves #gfp-1020), and Alexa Fluor 488/568 secondaries (ThermoFisher).

After immunolabeling, larvae were imaged on a LSM800 confocal microscope with a 63X/NA1.4 oil objective. Optimal z-sections were obtained to image the depth of the axon terminal. For quantification of immunofluorescence, a custom ImageJ macro was used to isolate the fluorescence signal from the GFP-labeled axons. A summed projection was generated and then the mean fluorescence intensity quantified. Mean fluorescence was normalized to background. Statistical analyses were done in JMP.

### In vivo analysis of endosome localization and transport in zebrafish

Larvae mosaically expressing endosome markers were generated as previously described (Drerup and Nechiporuk 2013). Briefly, zygotes were microinjected with DNA plasmids encoding a neuron specific promotor (*5kbneurod*) driving expression of the particular Rab fused to mRFP. Larvae expressing the construct of interest in pLL sensory neurons were isolated using an AxioZoom fluorescence dissecting microscope, mounted in 1.5% low melt agarose, and imaged on a LSM800 confocal microscope using a 63X/NA1.2 water immersion objective (Zeiss). For analyses of Rab5/7/11 vesicle localization, optimal z-stacks were obtained through the expressing pLL axon terminal. A standard deviation projection was then done in ImageJ and the total particle area quantified. The axon terminal area was also measured to allow a normalized endosome/axon terminal area quantification.

For analyses of vesicle transport, larvae were mounted similarly and imaged with the LSM800 confocal using the 63X/NA1.2 water immersion objective (Zeiss) as previously described (Drerup and Nechiporuk 2016). Images were taken 3 times per second for 150 seconds over a region of pLL axon in a single plane to track the location of moving puncta. Imaging sessions were analyzed using Kymograph analysis in Metamorph (Molecular Devices). Transport distance was quantified and statistical significance analyzed using JMP.

## REFERENCES

Aberle H, Haghighi AP, Fetter RD, McCabe BD, Magalhães TR, Goodman CS. 2002. *wishful thinking* encodes a BMP type II receptor that regulates synaptic growth in *Drosophila*. Neuron 33:545–58.

Bahloul A, Simmler MC, Michel V, Leibovici M, Perfettini I, Roux I, Weil D, Nouaille S, Zuo J, Zadro C, Licastro D, Gasparini P, Avan P, Hardelin JP, Petit C. 2009. Vezatin, an integral membrane protein of adherens junctions, is required for the sound resilience of cochlear hair cells. EMBO Mol Med 1:125–38.

Ball R W, Warren-Paquin, Tsurudome MK, Liao EH, Elazzouzi F, Cavanagh C, An BS, Wang TT, White JH, Haghighi AP. 2010 Retrograde BMP signaling controls synaptic growth at the NMJ by regulating Trio expression in motor neurons. Neuron 66: 536–49.

Burger A, Lindsay H, Felker A, Hess C, Anders C, Chiavacci E, Zaugg J, Weber LM, Catena R, Jinek M, Robinson MD, Mosimann C. 2016. Maximizing mutagenesis with solubilized CRISPR-Cas9 ribonucleoprotein complexes. Development 143:2025–37.

Carrington B, Varshney GK, Burgess SM, Sood R. 2015. CRISPR-STAT: an easy and reliable PCR-based method to evaluate target-specific sgRNA activity. Nucleic Acids Res 43:e157.

Danglot L, Freret T, Le Roux N, Narboux Nême N, Burgo A, Hyenne V, Roumier A, Contremoulins V, Dauphin F, Bizot JC, Vodjdani G, Gaspar P, Boulouard M, Poncer JC, Galli T, Simmler MC. 2012. Vezatin is essential for dendritic spine morphogenesis and functional synaptic maturation. J Neurosci 32:9007–22.

Deshpande M, Rodal AA. 2016. The Crossroads of Synaptic Growth Signaling, Membrane Traffic and Neurological Disease: Insights from *Drosophila*.Traffic 17:87–101.

Drerup CM, Nechiporuk AV. 2013. JNK-interacting protein 3 mediates the retrograde transport of activated c-Jun N-terminal kinase and lysosomes. PLoS Genet 9:e1003303.

Drerup CM, Nechiporuk AV. 2016. In vivo analysis of axonal transport in zebrafish. Methods Cell Biol 131:311–29.

Drerup CM, Herbert AL, Monk KR, Nechiporuk AV. 2017. Regulation of mitochondria-dynactin interaction and mitochondrial retrograde transport in axons. Elife pii: e22234.

Fisher B, Weiszmann R, Frise E, Hammonds A, Tomancak P, Beaton A, Berman B, Quan E, Shu S, Lewis S, et al. 2012. BDGP insitu homepage. http://insitu.fruitfly.org/cgi-bin/ex/insitu.pl.

Graziadio L, Palumbo V, Cipressa F, Williams BC, Cenci G, Gatti M, Goldberg ML, Bonaccorsi S. 2018. Phenotypic characterization of *diamond (dind*), a *Drosophila* gene required for multiple aspects of cell division. Chromosoma127:489–504.

Guedes-Dias P, Holzbaur ELF. 2019. Axonal transport: Driving synaptic function. Science 366(6462). pii: eaaw9997.

Guo W, Stoklund Dittlau K, Van Den Bosch L. 2019. Axonal transport defects and neurodegeneration: Molecular mechanisms and therapeutic implications. Semin Cell Dev Biol. 18. pii: S1084-9521(18)30219-2.

Higashi-Kovtun ME, Mosca TJ, Dickman DK, Meinertzhagen IA, Schwarz TL. 2010. Importin-beta11 regulates synaptic phosphorylated mothers against decapentaplegic, and thereby influences synaptic development and function at the *Drosophila* neuromuscular junction. J Neurosci 30:5253–68.

Hyenne V, Louvet-Vallée S, El-Amraoui A, Petit C, Maro B, Simmler MC. 2005. Vezatin, a protein associated to adherens junctions, is required for mouse blastocyst morphogenesis. Dev Biol 287:180–91.

Hyenne V, Souilhol C, Cohen-Tannoudji M, Cereghini S, Petit C, Langa F, Maro B, Simmler MC. 2007. Conditional knock-out reveals that zygotic *vezatin*-null mouse embryos die at implantation. Mech Dev 124:449–62.

Kimmel CB, Ballard WW, Kimmel SR, Ullmann B, Schilling TF. 1995. Stages of embryonic development of the zebrafish. Dev Dyn 203:253–310.

Koppel N, Friese MB, Cardasis HL, Neubert TA, Burden SJ. 2019. Vezatin is required for the maturation of the neuromuscular synapse. Mol Biol Cell 30:2571–83.

Krämer H, Phistry M. 1999. Genetic analysis of *hook*, a gene required for endocytic trafficking in *Drosophila*. Genetics. 1999 151:675–84.

Küssel-Andermann P, El-Amraoui A, Safieddine S, Nouaille S, Perfettini I, Lecuit M, Cossart P, Wolfrum U, Petit C. 2000. Vezatin, a novel transmembrane protein, bridges myosin VIIA to the cadherin-catenins complex. EMBO J 19:6020–9.

Lloyd TE, Machamer J, O’Hara K, Kim JH, Collins SE, Wong MY, Sahin B, Imlach W, Yang Y, Levitan ES, McCabe BD, Kolodkin AL. 2012. The p150(Glued) CAP-Gly domain regulates initiation of retrograde transport at synaptic termini. Neuron 74:344–60.

Lee C H, Herman TG, Clandinin TR, Lee R, Zipursky SL. 2001. N-cadherin regulates target specificity in the *Drosophila* visual system. Neuron 30:437–50.

Lee T, Luo L. 1999. Mosaic analysis with a repressible cell marker for studies of gene function in neuronal morphogenesis. Neuron 22:451–61.

Marchesin V, Castro-Castro A, Lodillinsky C, Castagnino A, Cyrta J, Bonsang-Kitzis H, Fuhrmann L, Irondelle M, Infante E, Montagnac G, Reyal F, Vincent-Salomon A, Chavrier P. 2015. ARF6-JIP3/4 regulate endosomal tubules for MT1-MMP exocytosis in cancer invasion. J Cell Biol 211:339–58.

Marie B, Sweeney ST, Poskanzer KE, Roos J, Kelly RB, Davis GW. 2004. Dap160/intersectin scaffolds the periactive zone to achieve high-fidelity endocytosis and normal synaptic growth. Neuron 43:207–19.

Marqués G, Bao H, Haerry TE, Shimell MJ, Duchek P, Zhang B, O’Connor MB. 2002. The *Drosophila* BMP type II receptor Wishful Thinking regulates neuromuscular synapse morphology and function. Neuron 33:529–43.

Marrus SB, Portman SL, Allen MJ, Moffat KG, DiAntonio A. 2004. Differential localization of glutamate receptor subunits at the *Drosophila* neuromuscular junction. J Neurosci 24:1406–15.

McCabe BD, Hom S, Aberle H, Fetter RD, Marques G, Haerry TE, Wan H, O’Connor MB, Goodman CS, Haghighi AP. 2004. Highwire regulates presynaptic BMP signaling essential for synaptic growth. Neuron 41:891–905.

McCabe BD, Marqués G, Haghighi AP, Fetter RD, Crotty ML, Haerry TE, Goodman CS, O’Connor MB. 2003. The BMP homolog Gbb provides a retrograde signal that regulates synaptic growth at the *Drosophila* neuromuscular junction. Neuron 39:241–54.

McKenney RJ, Huynh W, Tanenbaum ME, Bhabha G, Vale RD. 2014. Activation of cytoplasmic dynein motility by dynactin-cargo adapter complexes. Science 345:337–41.

Montagnac G, Sibarita JB, Loubéry S, Daviet L, Romao M, Raposo G, Chavrier P. 2009. ARF6 Interacts with JIP4 to control a motor switch mechanism regulating endosome traffic in cytokinesis. Curr Biol 19:184–95.

Moughamian AJ, Holzbaur EL. 2012. Dynactin is required for transport initiation from the distal axon. Neuron 74:331–43.

Moughamian AJ, Osborn GE, Lazarus JE, Maday S, Holzbaur EL. 2013. Ordered recruitment of dynactin to the microtubule plus-end is required for efficient initiation of retrograde axonal transport. J Neurosci 33:13190–203.

Narayanan R, Krämer H, Ramaswami M. 2000. *Drosophila* endosomal proteins hook and deep orange regulate synapse size but not synaptic vesicle recycling. J Neurobiol 45:105–19.

Neisch AL, Avery AW, Machamer JB, Li MG, Hays TS 2016. Methods to identify and analyze gene products involved in neuronal intracellular transport using *Drosophila*. Methods Cell Biol 131:277–309.

Obholzer N, Wolfson S, Trapani JG, Mo W, Nechiporuk A, Busch-Nentwich E, Seiler C, Sidi S, Söllner C, Duncan RN, Boehland A, Nicolson T. 2008. Vesicular glutamate transporter 3 is required for synaptic transmission in zebrafish hair cells. J Neurosci 28:2110–8.

O’Connor-Giles KM, Ho LL, Ganetzky B. 2008. Nervous wreck interacts with thickveins and the endocytic machinery to attenuate retrograde BMP signaling during synaptic growth. Neuron 58:507–18.

Persson U, Izumi H, Souchelnytskyi S, Itoh S, Grimsby S, Engström U, Heldin C H, Funa K, ten Dijke P. 1998. The L45 loop in type I receptors for TGF-beta family members is a critical determinant in specifying Smad isoform activation. FEBS Lett 434:83–87.

Rawson JM, Lee M, Kennedy EL, Selleck SB. 2003. *Drosophila* neuromuscular synapse assembly and function require the TGF-beta type I receptor saxophone and the transcription factor Mad. J Neurobiol 55:134–50.

Reck-Peterson SL, Redwine WB, Vale RD, Carter AP. 2018. The cytoplasmic dynein transport machinery and its many cargoes. Nat Rev Mol Cell Biol 19:382–398.

Roy S, Huang H, Liu S, Kornberg TB. 2014. Cytoneme-mediated contact-dependent transport of the *Drosophila* decapentaplegic signaling protein. Science 343:1244624.

Sanda M, Ohara N, Kamata A, Hara Y, Tamaki H, Sukegawa J, Yanagisawa T, Fukunaga K, Kondo H, Sakagami H. 2010. Vezatin, a potential target for ADP-ribosylation factor 6, regulates the dendritic formation of hippocampal neurons. Neurosci Res 67:126–36.

Schindelin J, Arganda-Carreras I, Frise E, Kaynig V, Longair M, Pietzsch T, Preibisch S, Rueden C, Saalfeld S, Schmid B, Tinevez JY, White DJ, Hartenstein V, Eliceiri K, Tomancak P, Cardona A. 2012. Fiji: an open-source platform for biological-image analysis. Nat Methods 9:676–82.

Schlager MA, Hoang HT, Urnavicius L, Bullock SL, Carter AP. 2014. In vitro reconstitution of a highly processive recombinant human dynein complex. EMBO J 33:1855–68.

Smith RB, Machamer JB, Kim NC, Hays TS, Marqués G. 2012. Relay of retrograde synaptogenic signals through axonal transport of BMP receptors. J Cell Sci 125:3752–64.

Sulkowski MJ, Han TH, Ott C, Wang Q, Verheyen EM, Lippincott-Schwartz J, Serpe M. 2016. A Novel, Noncanonical BMP Pathway Modulates Synapse Maturation at the *Drosophila* Neuromuscular Junction. PLoS Genet 12:e1005810.

Sunio A, Metcalf AB, Krämer H. 1999. Genetic dissection of endocytic trafficking in *Drosophila* using a horseradish peroxidase-bride of sevenless chimera: hook is required for normal maturation of multivesicular endosomes. Mol Biol Cell 10:847–59.

Szatmári Z, Kis V, Lippai M, Hegedus K, Faragó T, Lorincz P, Tanaka T, Juhász G, Sass M. 2014. Rab11 facilitates cross-talk between autophagy and endosomal pathway through regulation of Hook localization. Mol Biol Cell 25:522–31.

Urnavicius L, Lau CK, Elshenawy MM, Morales-Rios E, Motz C, Yildiz A, Carter AP. 2018. Cryo-EM shows how dynactin recruits two dyneins for faster movement. Nature 554:202–206.

Westerfield M. 1993. The Zebrafish Book: A Guide for the Laboratory Use of Zebrafish (Brachydanio rerio). Eugene: University of Oregon Press.

Williamson CD, Donaldson JG. 2019. Arf6, JIP3, and dynein shape and mediate macropinocytosis. Mol Biol Cell 30:1477–1489.

Wong MY, Zhou C, Shakiryanova D, Lloyd TE, Deitcher DL, Levitan ES. 2012. Neuropeptide delivery to synapses by long-range vesicle circulation and sporadic capture. Cell 148:1029–38.

Xiang X, Qiu R, Yao X, Arst HN Jr, Peñalva MA, Zhang J. 2015. Cytoplasmic dynein and early endosome transport. Cell Mol Life Sci 72:3267–80.

Yao X, Arst HN Jr, Wang X, Xiang X. 2015. Discovery of a vezatin-like protein for dynein-mediated early endosome transport. Mol Biol Cell 26:3816–27.

