## Supplemental figures 1-3 for "Vezatin is required for the retrograde axonal transport of endosomes in *Drosophila* and zebrafish"

**Supplemental Figure 1. Synaptic vesicles markers and an active zone marker localize normally in *vezl* mutants.**
**A-D,** Third-instar larval NMJs stained with anti-HRP (white); terminal boutons are to the right Scale bar is 10 μm. Unlike HRP, the SV proteins CSP (**A,B**) and Syt (**C,D**) are not enriched in terminal boutons in *vezl*$^{13.1}$*/Df(3R)Exel6180* mutants.
**E,F,** Structured illumination microscopy (SIM) images of terminal boutons bouton stained with antibodies against HRP (blue), the major AZ protein Brp (green) and the post-synaptic glutamate receptor subunit GluIIC (red). Scale bar is 10 μm. In both wild-type (**E**), Brp and GluIIC are in apposition. In *vezl*$^{13.1}$*/Df(3R)Exel6180* mutants (**F**)**,** Brp and GluIIC remain in apposition, but are at decreased density. Barren areas appear to correlate with areas having increased anti-HRP staining (merge panel has reduced HRP intensity to show detail).
**G,** Quantification of anti-Brp puncta density in terminal boutons. ***$p$<0.001, based on a two-tailed t-test.

**Supplemental Figure 2. Expression of a human Vezatin (hVez) transgene is unable to properly localize and cannot rescue loss of Vezl**
**A,** A wild-type third-instar larval NMJ at which human vezatin (hVez) tagged with Venus (green) is expressed in the motor neuron. Anti-HRP is in blue. Unlike Vezl:Venus, hVez:Venus is not enriched in the terminal bouton (top). Scale bar is 10 μm.
**B,** Quantification of NMJ bouton numbers. Expression of hVez:Venus in motor neurons does not restore NMJ size in *vezl*$^{13.1}$*/Df(3R)Exel6180* mutants or alter NMJ size in wild type. ***$p$<0.001, ns = not significant, based on two-tailed t-tests.

**Supplemental Figure 3. p150 accumulates in *vezt* mutant axon terminals**
**A-F**, Immunofluorescence analysis of the dynactin component p150, Dynein Heavy Chain (Dync1h1), and mitochondria (CytoC) in zebrafish pLL axon terminals marked by cytoplasmic GFP (green) in the *TgBAC(neurod:egfp)*$^{nl1}$ transgenic line. Arrowheads point to axon terminal swellings in *vezt* mutants. Scale bar is 10 μm.
**G,** Quantification of **A-F**. Values were normalized to background. p150 but not Dync1h1 or CytoC accumulates significantly in *vezt* mutant terminals.

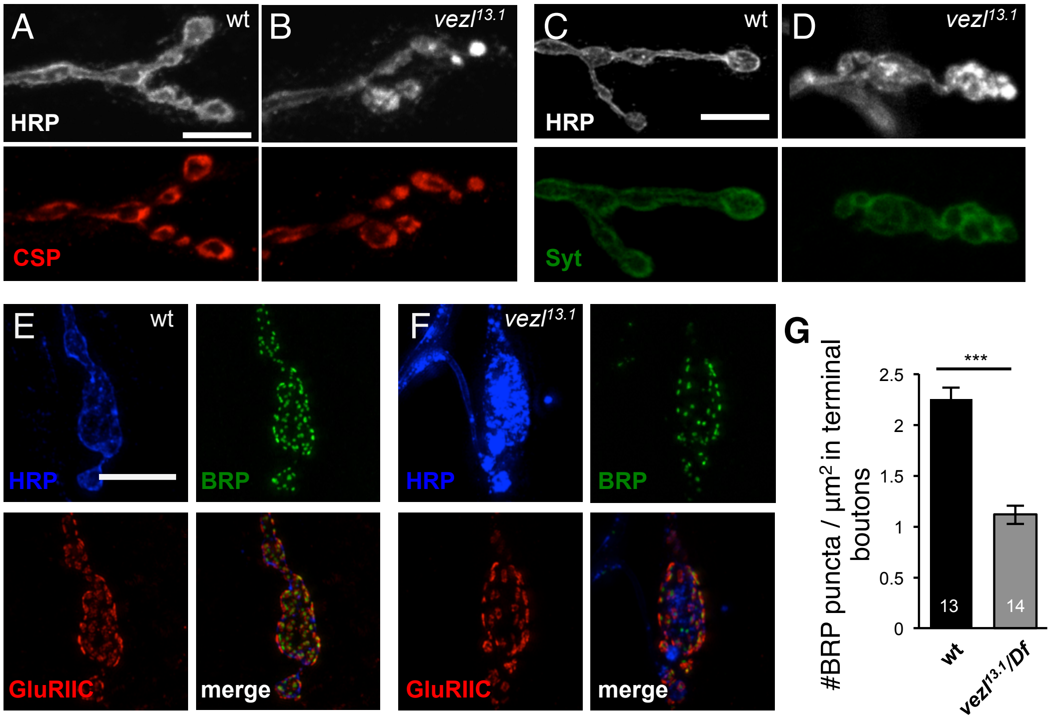


**Supp Figure 1**

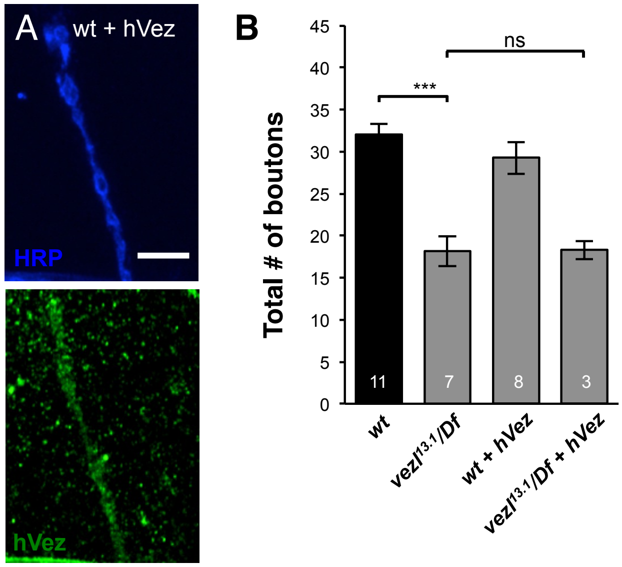


**Supp Figure 2**

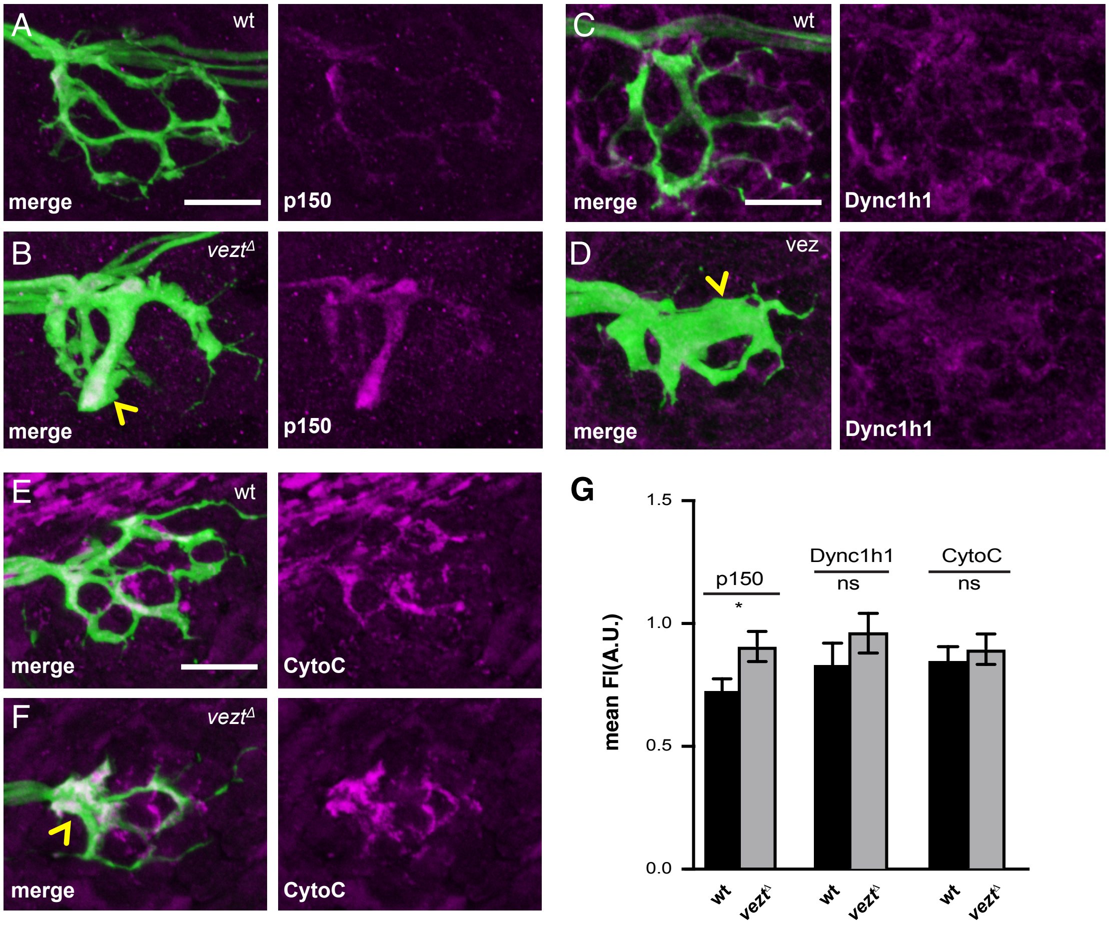


**Supp Figure 3**
